# Comparative Performance of the Finite Element Method and the Boundary Element Fast Multipole Method for Problems Mimicking Transcranial Magnetic Stimulation (TMS)

**DOI:** 10.1101/411082

**Authors:** Aung Thu Htet, Guilherme B. Saturnino, Edward H. Burnham, Gregory M. Noetscher, Aapo Nummenmaa, Sergey N. Makarov

## Abstract

A study pertinent to the numerical modeling of cortical neurostimulation is conducted in an effort to compare the performance of the finite element method (FEM) and an original formulation of the boundary element fast multipole method (BEM-FMM) at matched computational performance metrics. We consider two problems: (i) a canonic multi-sphere geometry and an external magnetic-dipole excitation where the analytical solution is available and; (ii) a problem with realistic head models excited by a realistic coil geometry. In the first case, the FEM algorithm tested is a fast open-source getDP solver running within the SimNIBS 2.1.1 environment. In the second case, a high-end commercial FEM software package ANSYS Maxwell 3D is used. The BEM-FMM method runs in the MATLAB^®^ 2018a environment.

In the first case, we observe that the BEM-FMM algorithm gives a smaller solution error for all mesh resolutions and runs significantly faster for high-resolution meshes when the number of triangular facets exceeds approximately 0.25 M. We present other relevant simulation results such as volumetric mesh generation times for the FEM, time necessary to compute the potential integrals for the BEM-FMM, and solution performance metrics for different hardware/operating system combinations. In the second case, we observe an excellent agreement for electric field distribution across different cranium compartments and, at the same time, a speed improvement of three orders of magnitude when the BEM-FMM algorithm used.

This study may provide a justification for anticipated use of the BEM-FMM algorithm for high-resolution realistic transcranial magnetic stimulation scenarios.

## 1. Introduction

For all three chief neurostimulation modalities – transcranial magnetic stimulation (TMS), transcranial electric stimulation (TES), and intracortical microstimulation (ICMS) – numerical computation of the electric fields within a patient-specific head model is the major and often only way to foster spatial targeting and obtain a quantitative measure of the required stimulation dose (Bikson et al., 2018). At present, a large portion of the macroscopic electromagnetic simulations of the brain are done using the finite element method (FEM). The FEM is widely used across engineering, physics, and geosciences. There are many general-purpose, open-source environments for FEM modeling, from high-level environments such as getDP (Dular et al., 1988), Deal.II (Bangerth et al., 2007), and FEniCS (Logg et al., 2012), to lower-level environments such as PETSc (Balay et al., 2018). Those solvers provide a practical and well-tested choice for creating problem-specific software solutions. Examples include:

- The well-known, open-source transcranial brain stimulation modeling software SimNIBS (Thielscher et al., 2015; Opitz et al., 2015; Nielsen et al., 2018), whose most recent version, v2.1, currently uses the open-source FEM software getDP (see Reference Manual 2017), which originates from the previous century (Dular et al., 1988);
- ROAST, a recently introduced TES modeling pipeline (Huang et al., 2018), which again uses the open-source FEM software getDP;
- COMETS: A MATLAB custom toolbox for simulating transcranial direct current stimulation (tDCS) (Jung et al., 2013; Lee et al., 2017), which is based on a stable and well-documented first-order FEM (Jin 2002).

On the other hand, there are proven and accurate commercial FEM solvers such as COMSOL Multiphysics and ANSYS Maxwell 3D, which are also able to accomplish the relevant simulation tasks.

In application to low-frequency bio-electromagnetic problems, the boundary element method (BEM) is also widely used, primarily for EGG/MEG modeling (Geselowitz 1967; Meijs et al., 1989; Hämäläinen et al., 1993; Ferguson et al., 1994; Mosher et al., 1999; Gramfort et al., 2014; Tadel et al., 2011; Gramfort et al., 2010; Stenroos et al., 2007; Stenroos and Sarvas 2012; Stenroos and Nummenmaa 2016; Nummenmaa et al 2013, Opitz et al., 2018; Rahmouni et al., 2018).

In application to high-frequency (or full-wave) electromagnetic problems solved via the surface/volume integral equation method, various accelerators have been proposed and employed, including the fast multipole method (FMM) (Song and Chew 1995; Song et al., 1997; Chew at al., 2001; Ergül and Gürel 2008), the fast Fourier transform (FFT) (Catedra 1995; Chen et al., 1996; Jin et al., 1996; Chen et al., 2004; Massey 2015), and the adaptive integral method (AIM) (Bleszynski et al., 1996; Wei and Yılmaz 2014; Massey et al., 2018).

However, FMM acceleration for low-frequency BEM brain modeling and in particular for TMS modeling, is much less common. The authors are aware of only one dedicated attempt to implement the FMM method, which was made over a decade ago (Kybic et al., 2005-2). In a recent study (Gomez et al., 2018), such a possibility was mentioned in the introduction but not immediately implemented. This in contrast to, for example, a low-frequency finite difference modeling technique, where a conceptually similar multi-grid formulation continues to demonstrate an impressive performance (Laakso and Hirata 2012; Laakso et al., 2018).

In a recent study (Makarov et al., 2018), the BEM-FMM approach was shown to be quite promising for TMS modeling and other relevant tasks. We have used the adjoint double-layer integral equation in terms of surface charge density (Rahmouni et al., 2018), pulse bases (piecewise-constant basis functions) with accurate integration of neighbor terms, simple Jacobi iterations, and an efficient and proven version of the FMM (Gimbutas and Greengard, 2015) originating from its inventors. The entire software package runs in the MATLAB environment.

In the present study, we further employ a generalized minimum residual (GMRES) iterative solution. We also convert a slower MATLAB loop, which corrects the FMM accuracy for neighboring facets via accurate integration, to a FORTRAN-based DLL. This increases the overall speed of the method more than a factor of two.

We next provide a detailed comparison of our method with the major “competitors” which are the above mentioned FEM-based TMS modeling tools. First, we compare the performance of the popular fast open-source FEM solver getDP within the SimNIBS 2.1.1 environment using a canonic multi-sphere problem where the analytical solution is available. Both the FEM software and the BEM-FMM software use matched performance metrics: the same multilayer sphere model, for which the analytical solution is available, the same server (Intel Xeon E5-26900 CPU 2.90 GHz), and the same operating system (Red Hat Enterprise Linux 7.5). Note that SimNIBS 2.1 and the BEM-FMM software support Linux and Windows operating systems. No effort to parallelize either of the methods (getDP FEM or BEM-FMM) has been made.

Second, we consider ten realistic CAD head models from the Population Head Model Repository (Lee et al., 2016; Lee et al., 2018; IT’IS Foundation 2016) augmented with a commercial TMS coil model and use a high-end commercial FEM software ANSYS Maxwell 18.2 2017 with adaptive mesh refinement and an arguably superior field accuracy. This study is a revision and extension of the comparison study started in (Makarov et al., 2018). Both the FEM and the BEM-FMM software use matched performance metrics: the same surface head model, the same coil model, the same server (Intel Xeon E5-2698 v4 CPU 2.2 GHz), and the same operating system (Windows Server 2016). Additionally, the high performance parallel computing (HPC) option within ANSYS with eight cores was used. No effort to parallelize BEM-FMM has been made. We compare field errors across all brain compartments and for every head model and establish the necessary computation times.

The study is organized as follows. Section 2 describes our BEM-FMM formulation in the general framework of the boundary element method for the Laplace equation. It also specifies integration of the fast multipole method and a correction approach for neighboring facets. Further, we describe two FEM solvers used for comparison and the corresponding comparison testbeds. Section 3 provides comparison results for the fast open-source FEM solver getDP in SimNIBS 2.1.1 environment, including both speed and relative accuracy versus the analytical solution. Section 4 provides method-to-method comparison results for realistic simulation scenarios, including use of the commercial FEM solver with adaptive mesh refinement, and establishes the level of agreement for volumetric field distribution. Section 5 discusses relevant aspects of both approaches (FEM versus BEM-FMM) and concludes the paper.

## 2. Materials and Methods

### 2.1. Boundary Element Method. Potential-based approach vs. charge-based approach

There exist two major types of the boundary integral equation for quasistatic modeling: the first is framed in terms of the electric potential, *φ*(***r***), while the second is written in terms of the electric charge density, *ρ*(***r***), at the boundaries (Barnard et al., 1967). The first is referred to as the double-layer formulation while the second one is known as the adjoint double-layer formulation (Rahmouni et al., 2018). Another or “symmetric” formulation also exists (Kybic et al., 2005-1, Rahmouni et al., 2018). The choice of the appropriate formulation depends on the problem under study. For TMS-related studies, we choose the adjoint double-layer formulation; this choice requires a more detailed explanation given below.

Consider two (or more) conducting compartments separated by some interface, *S*. The outer compartment has an electrical conductivity of *σ*_*out*_ and the inner compartment has electrical conductivity *σ*_*in*_ as shown in Fig. 1. The vector ***n***(***r***) in Fig. 1 is the outward unit normal vector for the inner compartment. When a given excitation electric field ***E***^*inc*^(***r***, *t*) is applied, surface electric charges with density *ρ*(***r***, *t*) will reside at *S* while the electric potential *φ*(***r***, *t*) and the normal component of the electric current density remain continuous across the interface. In the quasi-static or low-frequency approximation, the time dependence is purely parametric; it is therefore omitted after separation of variables.

**Fig. 1.**
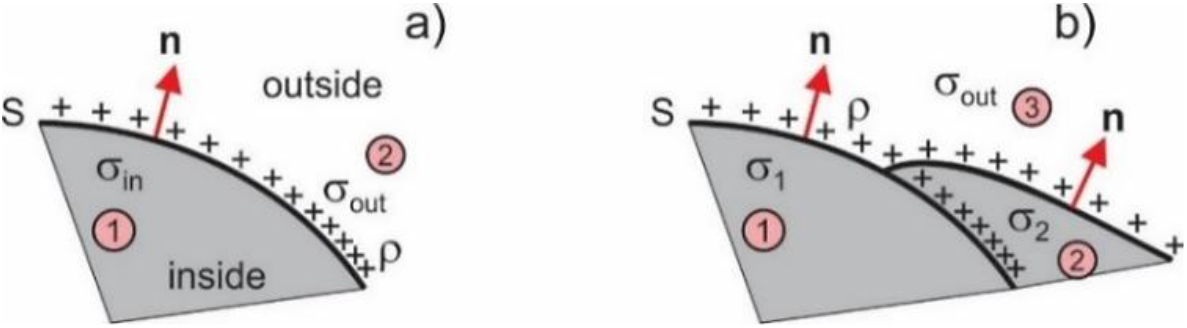
Boundary between two conducting compartments with different conductivities and surface charge density *ρ*(***r***) residing at the boundary.

### 2.2. Potential-based approach or double-layer formulation for EEG-MEG studies

The most widely used potential-based approach results in the integral equation (Barnard et al., 1967, Geselowitz 1967; Sarvas 1987, Meijs et al., 1989; Hämäläinen et al., 1993; Ferguson et al., 1994; Mosher et al., 1999; Stenroos et al., 2007)

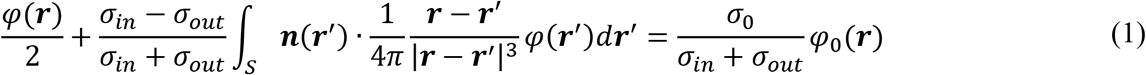

where *σ*_0_ = 1 S/m is the unit conductivity and *φ*_0_(***r***) is a given and conservative excitation. For EEG/MEG applications, it is an electric potential of a current dipole in an unbounded conducting medium with the unit conductivity *σ*_0_ = 1 S/m. Equation (1) is well suited for EEG studies since it gives the solution directly in the form of the electric potential (or voltage) on the scalp surface and on other interfaces. It is also well suited for MEG studies since the magnetic field is then straightforwardly found using Geselowitz’ formula (see, for example, Sarvas 1987).

At the same time, Eq. (1) has a few limitations. First, it is derived using Green’s second identity (Hämäläinen et al., 1993) and is therefore only valid for closed surfaces with one value of external conductivity, in particular for surfaces enclosed into each other in the form of an onion structure. Inclusion of surface junctions (e.g. an opening in the skull sketched in Fig. 1b) requires a special treatment (Stenroos, 2016). Next, the excitation in Eq. (1) must be a conservative field: the solenoidal-field excitation of a TMS coil is not allowed. In addition, the FMM would require computing the potential of a double layer as opposed to a single-layer gradient.

### 2.3. Modification of the potential-based approach for TMS studies

In order to overcome the limitation of the conservative excitation and use Eq. (1) for TMS studies, a reciprocity principle has been employed (Heller and van Hulsteyn, 1992, Nummenmaa et al., 2013). This principle allows us to reuse the standard MEG computational methods based on Eq. (1). Assume that the TMS coil is approximated by a number of equivalent time-varying magnetic dipoles, ***m***. Then, a total electric field ***E***^*t*^ induced by one such dipole ***m*** at a certain location ***r***_1_ within the brain compartments is computed via an external magnetic field generated by a reciprocal oscillating current dipole located at ***r***_1_. The last task is solved with Eq. (1) and Geselowitz’ formula. However, in order to find the vector electric field at a single location due to a single magnetic dipole, we have to solve Eq. (1) three times, which is less convenient.

### 2.4. Charge-based approach or adjoint double-layer formulation used in this study

The surface-charge formulation might have several advantages for the present and potentially other tasks. First, the excitation field does not have to be conservative. The corresponding integral equation is simply obtained by writing the total electric field ***E***^*t*^ in a form that takes into account the non-conservative external field ***E***^*inc*^ of the TMS coil(s) and a conservative contribution of the secondary induced surface charge density (time dependence is not shown)

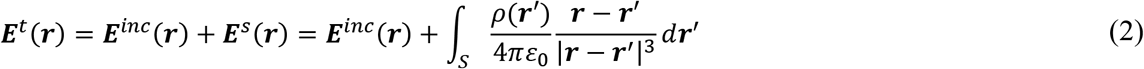

where *ε*_0_ is the electric permittivity of vacuum. Taking the limit of Eq. (2) as ***r*** approaches the surface *S* from both sides and using the continuity condition for the normal current component, *σ****E***^*t*^(***r***), one obtains the adjoint double-layer equation (Barnard et al., 1967, Makarov et al., 2016, Rahmouni et al., 2018, Makarov et al., 2018 (Supplement)) for the surface charge density in the following form:

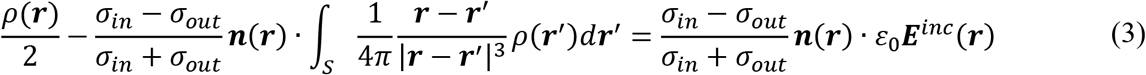

Note that the scaling constant *ε*_0_ is indeed redundant. Equation (3) is directly applicable to any excitation field, without using the reciprocity principle. Additionally, the surface junction case from Fig. 1b is permitted. In contrast to Eq. (1), the normal-vector multiplication becomes external in the adjoint operator. This leads to computing the gradient of a single layer, which, from the viewpoint of the FMM, might be more beneficial than computing the potential of a double layer for Eq. (1). Finally, the electric field distribution just inside/outside cortical surfaces, which is most important for TMS, is almost trivially computed from the already known charge solution:

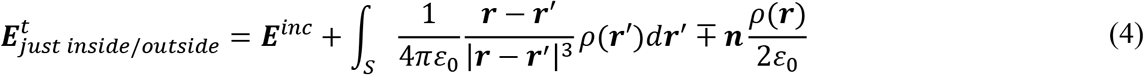

### 2.5. Fast Multipole Method (FMM)

The fast multipole method introduced by Rokhlin and Greengard (Rokhlin, 1985; Greengard and Rokhlin, 1987) speeds up computation of a matrix-vector product by many orders of magnitude. Such a matrix-vector product naturally appears when a an electric field from many point sources *ρ*(***r***^′^) in space has to be computed at many observation or target points ***r***. In other words, it is the discretization of the surface integral in Eq. (2) or in Eq. (3). Assuming piecewise constant expansion basis functions (pulse bases), one has

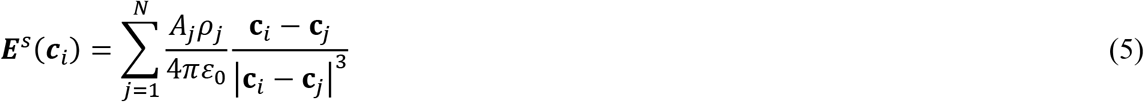

where *A*_*i*_, ***c***_*i*_, *i* = 1, …, *N* are, respectively, areas and centers of the triangular surface facets *t*_*i*_ while *ρ*_*i*_ are surface charge densities at the patch centers. Approximation (5) is computed via the FMM. We adopt, integrate, and use an efficient and proven version of the FMM (Gimbutas and Greengard, 2015) originating from its inventors. In this version, there is no a priori limit on the number of levels of the FMM tree, although after about thirty levels, there may be floating point issues (L. Greengard, private communication). The required number of levels is determined by a maximum permissible least-squares error or method tolerance, which is specified by the user. The FMM is a FORTAN 90/95 program compiled for MATLAB. The tolerance level `iprec` of the FMM algorithm is set at 0 (the relative least-squares error is guaranteed not to exceed 0.5%). This FMM version allows for a straightforward inclusion of a controlled number of analytical neighbor integrals to be precisely evaluated as specified below.

### 2.6. Correction of neighboring terms. Iterative solution

Approximation (5) is inaccurate for the neighbor facets. In the framework of Petrov-Galerkin method with the same pulse bases as testing functions, it is corrected as follows

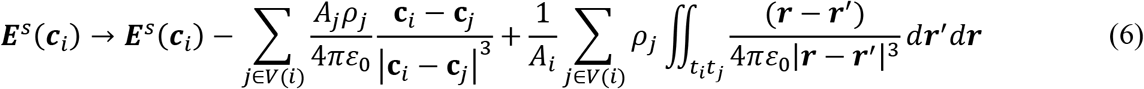

where *V*(*i*) is a neighborhood of observation triangle *t*_*i*_. Inner integrals in Eq. (6) are computed analytically (Wilton et al., 1984, Wang et al., 2003; Makarov et al., 2016); the outer integrals use a Gaussian quadrature of 10^th^ degree of accuracy (Cools, 2003). We have implemented two methods: a ball neighborhood with a radius *R* such that |**c**_*i*_ − **c**_*j*_| ≤ *R* and a neighborhood of *K* ≪ *N* nearest triangular facets. Both methods provide similar results. However, the second method leads to fixed-size arrays of precalculated double surface integrals and to a faster speed. Therefore, it has been preferred. Inclusion of a small number of precomputed neighbor integrals (three to twelve) drastically improves the convergence of the iterative solution; inclusion of a larger number has little, if any effect. The present BEM-FMM approach performs precise analytical integration over the 12 closest neighbor facets. For postprocessing computations of the volumetric E-field from the known charge distribution, analytical integration is desired very close to the boundaries.

Equation (3) is solved iteratively using the native MATLAB GMRES (generalized minimum residual method) of Drs. P. Quillen and Z. Hoffnung of MathWorks, Inc. Although this method may be somewhat slower than a simplified in-house version of the GMRES, its overall performance and convergence are excellent, especially for complicated head geometries. The relative residual of the BEM-FMM iterative method is set as 1e-4; the number of iterations does not exceed 30.

### 2.7. Finite Element Method: getDP solver used in SimNIBS 2.1.1

SimNIBS 2.1.1 employs the default FEM solver: the open source fast FEM software package called `getDP` (see Reference Manual, 2017). In SimNIBS, getDP is configured to use the PETSc conjugate gradient (CG) solver with a relative residual of 1e-9 and the incomplete Cholesky (ICC) preconditioner with 2 factor levels. After the FEM solution is completed, field interpolation for arbitrary points in space is accomplished using the “super-convergent approach” (or SCA) recently implemented in SimNIBS 2.1.1. In this approach, the original tessellation is preserved, and the electric field at the nodes is interpolated from the electric field values at the tetrahedra centers. Following this, further linear interpolation for arbitrary observation points is performed using the original tessellation (Zienkiewicz and Zhu, 1992). To avoid problems due to discontinuities of the electric field across tissue boundaries, the field recovery is performed for each brain compartment separately.

### 2.8. Hardware

All comparison results reported in the following section and related to the multi-sphere solution are obtained using the same server and operating system: an Intel Xeon E5-2690 CPU at 2.90 GHz with 192 Gbytes of RAM running Red Hat Enterprise Linux 7.5. We run MATLAB version 2018a in Linux.

No effort to parallelize either of the methods (getDP FEM or BEM-FMM) has been made. However, both used software packages – getDP and core MATLAB – by default perform multithreading pertinent to linear algebra operations available in LAPACK (Linear Algebra PACKage) and some level-3 BLAS (Basic Linear Algebra Subprograms) matrix operations, allowing them to execute faster on multicore-enabled machines.

### 2.9 Comparison testbed for multi-sphere solution

The comparisons are carried out in models consisting of four-layered spheres, as adopted from Engwer et al., 2017 and Piastra et al., 2018. Although both of these references are concerned with MEG and EEG dipoles, the corresponding models are equally applicable to the present problem, which is also closely related to the MEG problem (Sarvas, 1987).

Figure 2a shows the problem geometry. The conductivity values are consistent with Engwer et al., 2017 and Piastra et al., 2018. To assure the test-grade surface triangulation, we first create *six* individual high-quality triangular base sphere meshes with the number of triangular facets ranging from approximately 0.011 M to 0.411 M (from lower to higher mesh density), using a high-quality surface mesh generator (Persson 2005; Persson and Strang, 2004), and implemented in MATLAB^®^. The minimum triangle mesh quality (twice the ratio of inscribed to circumscribed circle radii for a triangular facet) is no less than 0.7, so that all the triangular facets are nearly equilateral. All triangles have nearly the same size.

**Fig. 2.**
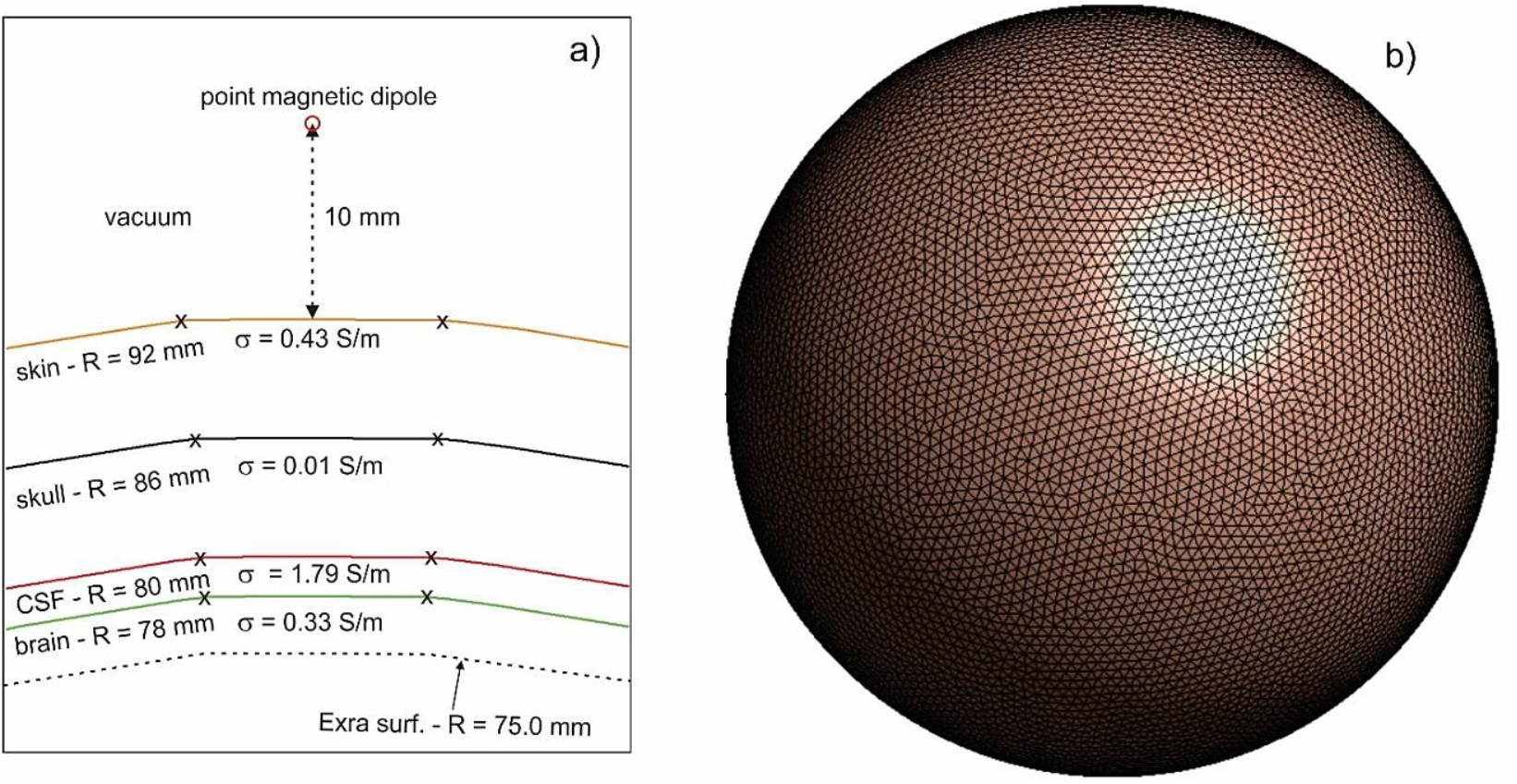
a) – Model geometry; b) – surface mesh topology for sphere #2 with the mesh resolution of 2.9 mm and the mesh density of 0.14 nodes/mm^2^.

Following this, we create six respective multi-sphere models by cloning and scaling every individual sphere mesh four times, as required by Fig. 2a. These “onion” models will be labeled #1 through #6. Additionally, every triangulated subsurface is also slightly scaled outwards so that its total area is exactly the sphere area with the prescribed radius. Table 1 lists the corresponding surface mesh resolution (or model resolution) and the mesh density (number of nodes per unit area) in the set of models. The mesh resolution is defined as the average edge length. The mesh density is given in nodes/mm^2^, which is a common measure in SimNIBS.

**Table. 1.** Model resolution and mesh density in every four-layer sphere model with a dummy sphere (see below).

| Model # | Facets total | Mesh resolution, mm | Mesh density, nodes/mm<sup>2</sup> |
| --- | --- | --- | --- |
| 1 | 0.06 M | 4.2 | 0.07 |
| 2 | 0.12 M | 2.9 | 0.14 |
| 3 | 0.24 M | 2.0 | 0.28 |
| 4 | 0.47 M | 1.4 | 0.55 |
| 5 | 1.03 M | 0.98 | 1.21 |
| 6 | 2.06 M | 0.69 | 2.41 |

The field error specified by Eq. (11) below is measured on two observation sphere surfaces. One of them is located 0.5 mm below the brain surface in Fig. 2a and has the radius of 77.5 mm. Another is located 1.5 mm below the brain surface in Fig. 2a and has the radius of 76.5 mm. Note that the FEM may have an insufficient resolution in regions where volumetric mesh density is low. In order to provide a fair comparison and assure the proper and sufficient FEM volumetric meshing in this observation domain, we introduce a fifth sphere with the radius of 75 mm into the model as shown in Fig. 2a. This sphere is a dummy object: its conductivity is equal to the brain conductivity of 0.33 S/m, so that the corresponding conductivity contrast is equal to zero. However, this dummy sphere is explicitly present in the FEM discretization. The mesh size of the combined model (four nontrivial brain compartments plus one dummy sphere) ranges from 0.06 M to 2.06 M facets in Table 1.

The excitation is given by a point magnetic dipole (a small loop of current), schematically shown in Fig. 2a, and located 10 mm above the skin surface. A magnetic dipole with the moment ***m***(*t*) located at point ***r***_2_ generates the magnetic vector potential given by,

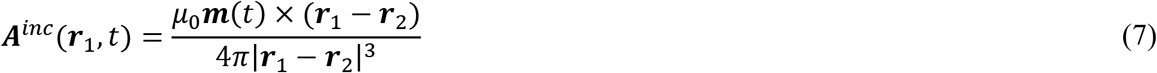

where ***r***_1_ is an arbitrary observation point and *μ*_0_ is the magnetic permeability of vacuum. From Eq. (7), the solenoidal electric field of the dipole in free space becomes

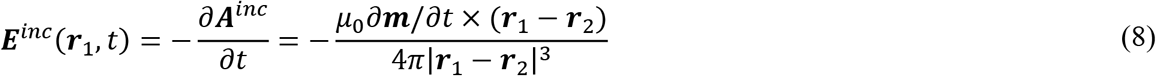

Further, we assume harmonic excitation of the form ***m***(*t*) = ***m***_0_exp (+*jωt*), convert to phasors, and eliminate the redundant constant phase factor of *j* using multiplication by *j*. This gives us the “static” real-valued excitation field

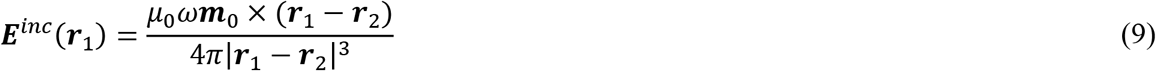

which could indeed be treated as a result of the separation of the time dependence and the spatial dependence, respectively.

For a dipole outside a spherical model with a spherically-symmetric conductivity distribution, the corresponding analytical solution neither depends on the individual sphere radii nor on the specific conductivity values (Sarvas 1987). The same manipulations that lead to Eq. (9) allow us to obtain from Ref. Sarvas 1987 an expression for the total field ***E*** in the form:

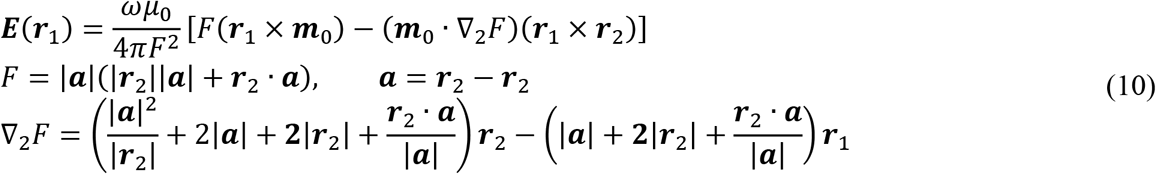

where ***r***_1_ is now an arbitrary observation point *within* the sphere model.

Once the analytical and numerical solutions is available, we compute the relative vector electric-field error using a matrix norm, that is

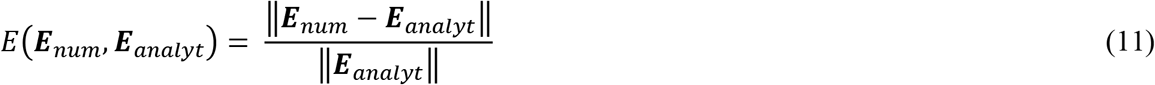

where 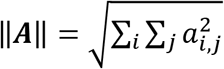 is the Frobenius or the *L*_2,2_ norm and ***E***_*num*_ ***E***_*analyt*_ are *M* × 3 electric field vector arrays at *M* observation points obtained with the numerical and analytical solutions, respectively. In all 6 models, we observed the vector electric field at *M*= 47,500 triangle centers of an observation sphere, with the radius of either 77.5 mm (0.5 mm below the “brain surface”) or 76.5 mm (further apart or 1.5 mm below the “brain surface”), respectively. This observation sphere has the mesh resolution of 2 mm and the mesh density of 0.3 nodes/mm^2^; it was generated as explained above. Since all the facets of the observation sphere have approximately the same area, the correction of Eq. (11) by a point-by-point area multiplication is insignificant.

### 2.10 Comparison testbed for realistic head models

Along with the standard multi-sphere solution described above, a population-based study has been performed to establish BEM-FMM accuracy for a more realistic TMS scenario. Ten high-resolution head models from the Population Head Model Repository (Lee et al, 2016; Lee et al., 2018; IT’IS Foundation, 2016) based on Connectome Project data (Van Essen et al, 2012) have been considered and augmented with the following material conductivities: scalp – 0.333 S/m, skull – 0.0203 S/m, CSF – 2.0 S/m, GM – 0.106 S/m, cerebellum – 0.126 S/m, WM – 0.065 S/m, ventricles – 2.0 S/m. Each head model has approximately 0.7 M facets in total. The mesh density in nodes/mm^2^ for each cavity is 0.8 (skin), 1.4 (skull), 4.9 (CSF), 3.7 (GM), 3.8 (WM), and 9.5 (ventricles).

For each head model, simulations with our method and simulations with the high-end commercial FEM software Maxwell 3D of ANSYS^®^ Electronics Desktop 2017 2.0, Release 18.2.0 have been performed; the FEM software used adaptively refined tetrahedral meshes. This study is a revision and extension of the corresponding comparison study started in (Makarov et al., 2018). The FEM software employs a T-Ω formulation with Ω being the nodal-based magnetic scalar potential, defined in the entire solution domain, and T being the edge-based electrical vector potential, defined only in the conducting eddy-current region. A Maxwell 3D project with Neumann boundary conditions, 4-5 adaptive mesh refinement passes, 30% mesh refinement rate per pass resulting in a final FEM mesh with 4-8 M tetrahedra, and a global energy error below 0.1% was employed.

Both the FEM software and the BEM-FMM software now use matched performance metrics and geometries: the same surface head and coil models, the same server (Intel Xeon E5-2698 v4 CPU 2.2 GHz with 256 Gbytes of RAM), and the same operating system (Windows Server 2016). Additionally, the high performance parallel computing (HPC) option in ANSYS was used with eight cores.

Three components of the electric field within the head along an observation line have been found using the two methods and were compared to each other. This observation line coincides with the centerline of the coil (figure-of-eight coil MRi-B91, MagVenture), which is located 9 mm above the approximate vertex location for each head. The coil is modeled in the form of solid conductors in the ANSYS Maxwell FEM software and in the form of a large number of elementary current sources (straight wire segments) in BEM-FMM, as shown in Fig. 3. The number of segments in our coil model is about 0.1 M; the BEM-FMM performance is not significantly affected due to the high speed of the FMM. Coil current is 5 kA and the excitation frequency is 3 kHz.

**Fig. 3.**
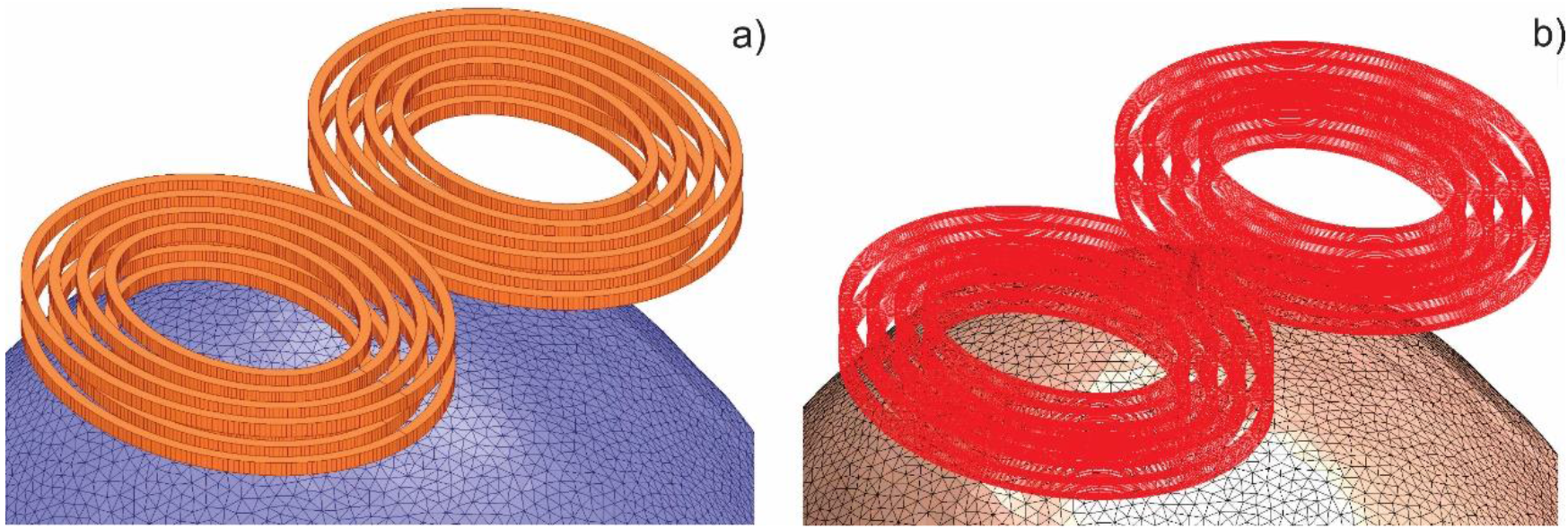
a) – Solid-conductor coil model in ANSYS FEM software and; b) – wire-conductor coil model in BEMFMM software.

## 3. Results for the multi-sphere solution. Comparison with getDP solver

### 3.1 Performance of getDP solver within the SimNIBS 2.1.1 environment

Table 2 presents run times for the FEM solution and the corresponding relative error Eq. (11) in the electric field computations, respectively. We consider two distinct observation spheres located 0.5 mm and 1.5 mm beneath the brain surface in Fig. 2a. The `getDP` FEM software was unable to process the largest problem with 2.06 M triangles on the server used in this study.

**Table. 2.** Speed and accuracy of the `getDP` solver within the SimNIBS 2.1.1 environment. The observation sphere is located 0.5 mm and 1.5 mm below the brain surface in Fig. 2a and has the radius of either 77.5 mm or 76.5 mm.

| Model # | Facets total | FEM solution time, sec | FEM E-field error (using SCA) 0.5 mm below the brain surface, % | FEM E-field error (using SCA) 1.5 mm below the brain surface, % |
| --- | --- | --- | --- | --- |
| 1 | 0.06 M | 8 | 6.5 | 2.9 |
| 2 | 0.12 M | 34 | 6.1 | 2.5 |
| 3 | 0.24 M | 95 | 5.8 | 2.0 |
| 4 | 0.47 M | 458 | 3.0 | 0.6 |
| 5 | 1.03 M | 1766 | 2.0 | 0.4 |
| 6 | 2.06 M | NA | NA | NA |

### 3.2 Performance of BEM-FMM solver within the MATLAB 2018a Linux environment

Table 3 presents run times for the BEM-FMM solution and the corresponding relative error Eq. (11) in the electric field computations, respectively. We again consider two distinct observation spheres located 0.5 mm and 1.5 mm beneath the brain surface in Fig. 2a. The tolerance level `iprec` of the FMM algorithm is set at 0 (the relative least-squares error is guaranteed not to exceed 0.5%). The relative residual of the BEM-FMM iterative method is set as 1e-4.

**Table. 3.** Speed and accuracy of the BEM-FMM solver within MATLAB 2018a Linux environment. The observation sphere is located 0.5 mm and 1.5 mm below the brain surface in Fig. 2a.

| Model # | Facets total | BEM-FMM solution time, sec | BEM-FMM E-field error 0.5 mm below the brain surface, % | BEM-FMM E-field error 1.5 mm below the brain surface, % |
| --- | --- | --- | --- | --- |
| 1 | 0.06 M | 9 | 2.3 | 2.4 |
| 2 | 0.12 M | 19 | 1.5 | 1.5 |
| 3 | 0.24 M | 34 | 0.6 | 0.7 |
| 4 | 0.47 M | 73 | 0.6 | 0.3 |
| 5 | 1.03 M | 151 | 0.2 | 0.2 |
| 6 | 2.06 M | 259 | 0.2 | 0.2 |

### 3.3 Comparison of pre-processing effort

The FEM approach requires volumetric tetrahedral mesh generation based on the CAD surface model. This operation has to be completed only once for each model, but it may require a significant amount of time. Table 4 reports volumetric mesh generation times in SimNIBS 2.1.1 for the six multi-sphere models. The mesh generation process is not parallelized. In SimNIBS 2.1.1, the volume meshing is performed using Gmsh (Geuzaine et al., 2009) using the frontal algorithm implemented in Tetgen (Si, 2015).

**Table. 4.** Volumetric (tetrahedral) mesh generation in SimNIBS 2.1.1.

| Model # | Facets total | Tetrahedra total | Meshing time, sec |
| --- | --- | --- | --- |
| 1 | 0.06 M | 0.26M | 7 |
| 2 | 0.12 M | 0.72M | 21 |
| 3 | 0.24 M | 1.8M | 66 |
| 4 | 0.47 M | 5.05M | 219 |
| 5 | 1.03 M | 16.43M | 988 |
| 6 | 2.06 M | 44.64M | 2678 |

While the FEM requires volumetric mesh generation, the BEM-FMM requires precomputing and storing potential integrals for the neighbor triangles. The number of neighbors is typically 3-12. This operation must be completed only once for each model. It is based on a `for`-loop over all triangular facets and is trivially parallelizable in MATLAB using the `parfor` syntax. Table 5 reports execution times for potential-integral computations and writing data to file, given 12 triangular neighbor facets and using the `parfor`-loop with 16 cores (`parpool(16)`) in MATLAB.

**Table. 5.** Times necessary for precomputing and storing potential integrals for the BEM-FMM algorithm given 12 neighbors and using the the `parfor`-loop with 16 cores (`parpool(16)`) in MATLAB.

| Model # | Facets total | Computation time for integrals, sec | Saving *.mat file, sec |
| --- | --- | --- | --- |
| 1 | 0.06 M | 35 | 1 |
| 2 | 0.12 M | 42 | 2 |
| 3 | 0.24 M | 55 | 3 |
| 4 | 0.47 M | 82 | 6 |
| 5 | 1.03 M | 143 | 11 |
| 6 | 2.06 M | 253 | 19 |

### 3.4 Comparison of post-processing effort – restoration of electric field at a surface

Here, we compare the speed of the super-convergent approach (SCA) recently implemented in SimNIBS 2.1.1 and the BEM-FMM field restoration algorithm for the observation sphere located at 1.5 mm below the brain surface in Fig. 2a. Table 6 presents the corresponding run-times. Since the potential integrals are not computed at this stage, the BEM-FEM is reduced to the plain FMM and is therefore very fast. If the potential field integrals were included, the post-processing time would increase by about 1 min without a significant effect on the solution accuracy. We also emphasize that the SCA algorithm is not yet optimized, and that it is possible to construct an interpolant, and store it to very significantly speed up future computations.

**Table. 6.**
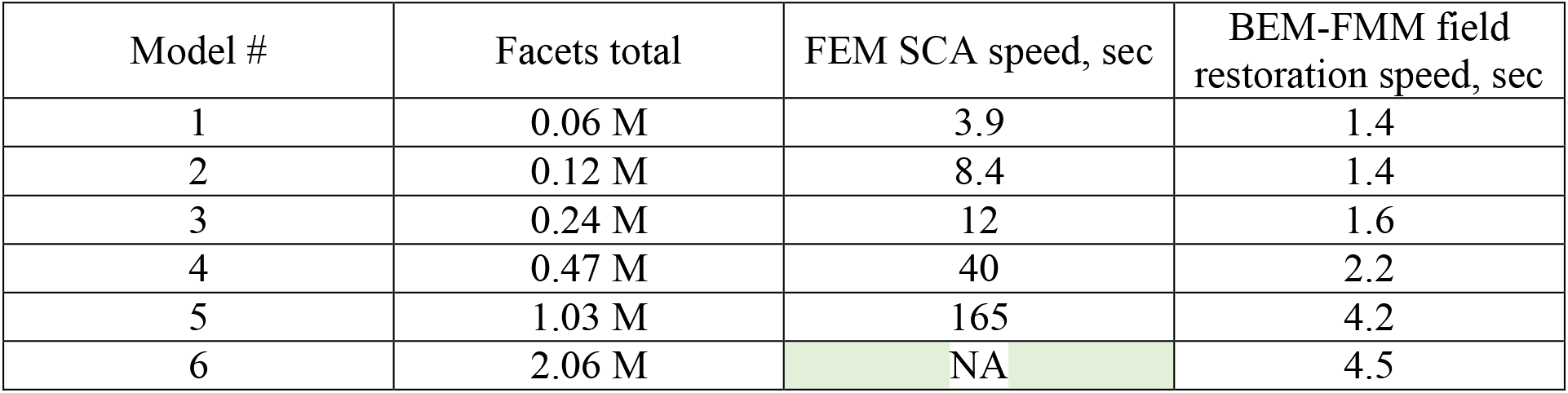
Post-processing run times of the SCA and the BEM-FEM engine when the observation sphere is located at 1.5 mm below the brain surface in Fig. 2a.

| Model # | Facets total | FEM SCA speed, sec | BEM-FMM field restoration speed, sec |
| --- | --- | --- | --- |
| 1 | 0.06 M | 3.9 | 1.4 |
| 2 | 0.12 M | 8.4 | 1.4 |
| 3 | 0.24 M | 12 | 1.6 |
| 4 | 0.47 M | 40 | 2.2 |
| 5 | 1.03 M | 165 | 4.2 |
| 6 | 2.06 M | NA | 4.5 |

### 3.5. Operating system and hardware performance for the BEM-FMM engine

We have also compared the BEM-FMM solution times using Linux- and Windows-based machines:

A. Intel Xeon E5-2690 CPU at 2.9 GHz, Red Hat Enterprise Linux 7.5; MATLAB 2018a Linux;
B. Intel Xeon E5-2698 v4 CPU at 2.2 GHz, Windows Server 2016; MATLAB 2018a Windows. Surprisingly, Server B significantly outperforms Server A, most likely due to the increased L1 cache size and multithreading capabilities of the E5-2698 vs. the E5-2690.

### 3.6. Summary of major comparison results for the multi-layered sphere

Figure 4 below summarizes results from Tables 2 and 3 for both methods and for the matched computational performance metrics. For the FEM solution, the superconvergent interpolation is used. For the BEM-FMM solution, the prescribed value of the relative residual is equal to 1e-4.

**Fig. 4.**
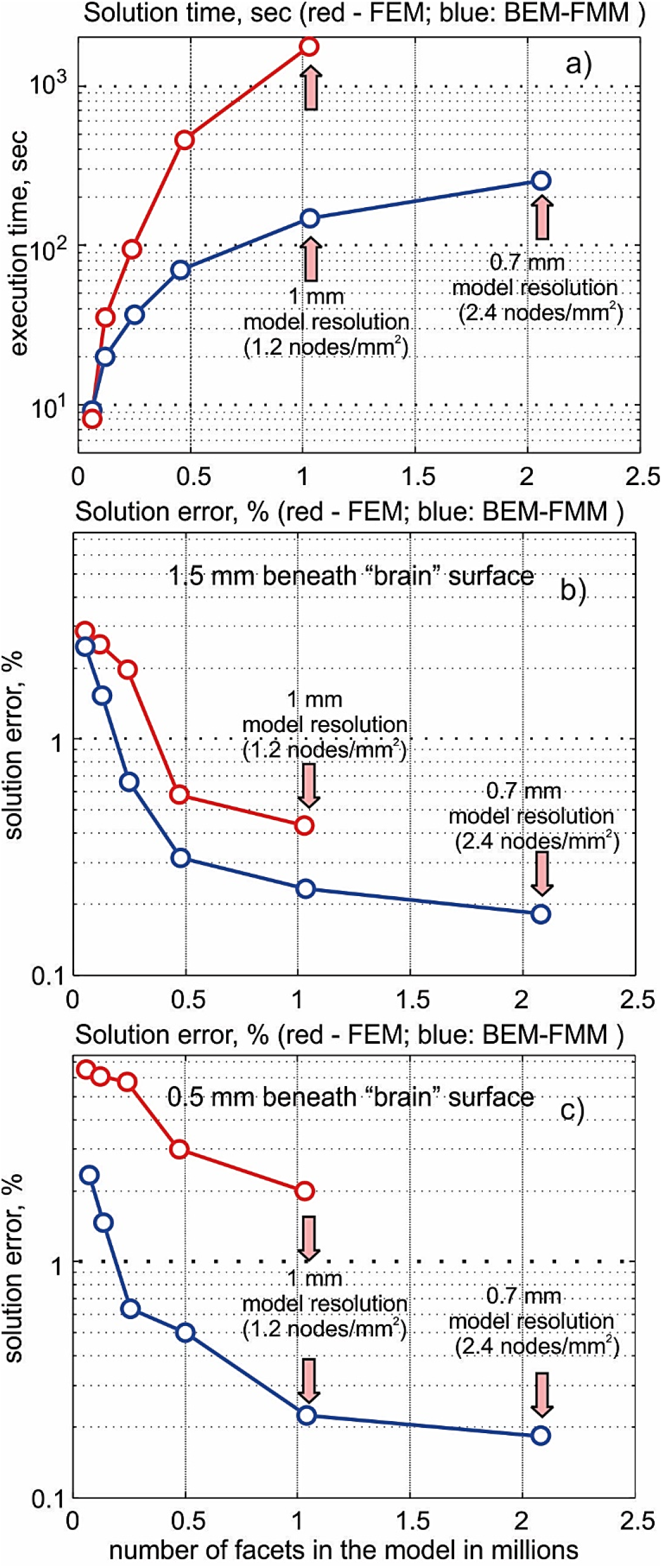
a) – Solution time; b) – solution error of the FEM and BEM-FMM algorithms, respectively, both as functions of the number of facets in the model (model resolution and/or mesh density) at 1.5 mm beneath the “brain” surface; c) – the same result at 0.5 mm beneath the “brain” surface.

Figure 4a shows the corresponding simulation times of the main algorithm. The BEM-FMM algorithm begins to outperform the FEM algorithm when the number of surface facets exceeds approximately 100,000. We observe that the BEM-FMM method runs much faster for high-resolution models. Figure 4b shows the relative error in the electric field Eq. (11) for the observation surface located 1.5 mm under the “brain” surface in Fig. 2a. We observe that the BEM-FMM method gives a smaller solution error for all mesh resolutions. The result does not change significantly when the observation surface is moved farther away from the brain interface. Figure 4c shows the relative error in the electric field Eq. (11) for the observation surface located 0.5 mm under the “brain” surface in Fig. 2a. We observe that the BEM-FMM method gives a much smaller solution error for all mesh resolutions.

The speed advantage of the BEM-FMM algorithm also holds for the pre- and post-processing steps as evidenced by Tables 5 and 6. The speed of the BEM-FMM algorithm can further be improved by switching from default complex arithmetic to real arithmetic (resulting in an increase by a factor of two).

## 4. Results for ten realistic head models. Comparison with commercial FEM solver ANSYS Maxwell 3D for intracranial fields

Figure 5 shows the computation geometry including the observation line and the representative comparison results for head #101309 (the first head model). This figure also illustrates the field distribution along the coil axis: the largest primary component *E*_*y*_ (Fig. 5a), the secondary yet somewhat significant component *E*_*z*_ (Fig. 5b), and the vanishingly small secondary component *E*_*x*_ (Fig. 5b as well). In Fig. 5, coil current is 5 kA and the excitation frequency is 3 kHz. All contour plot values are in V/m.

**Fig. 5.**
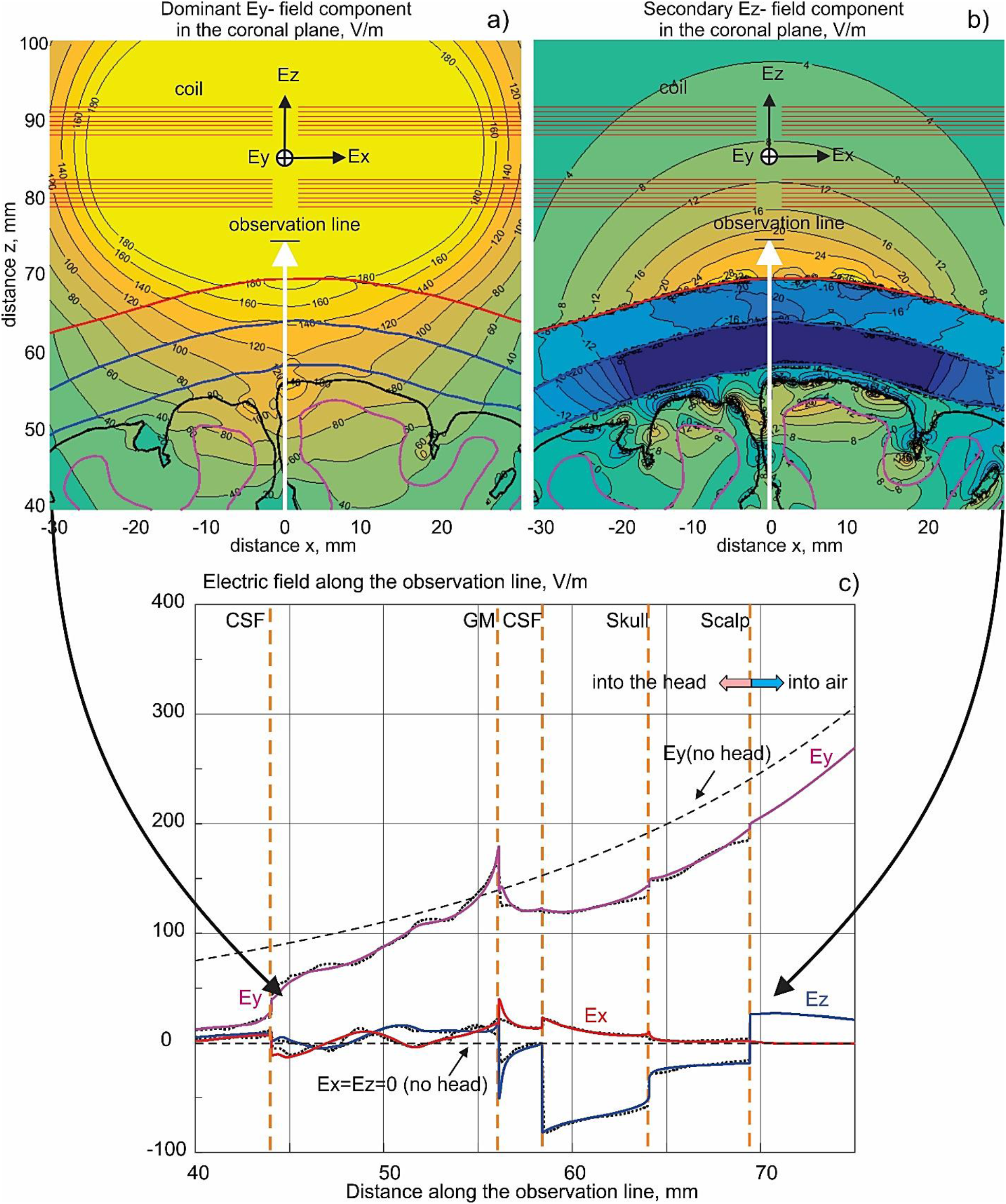
a, b) – Computation geometry, position of the observation line, and surface field distributions for head #101309 given 5 kA of coil current at 3 kHz; c) – electric field comparison along the line. BEM-FMM solution is shown by solid curves; the ANSYS Maxwell 3D FEM solution is given by dotted curves.

The Maxwell 3D project with Neumann boundary conditions, 4 adaptive mesh refinement passes, 30% mesh refinement rate per pass resulting in the final FEM mesh with approximately 5 M tetrahedra, and a global energy error below 0.1% was employed. The BEM-FMM solution uses 20 iterations (relative residual is below 0.1% and execution time is about 100 sec on the 2.2 GHz server), the analytical integration with twelve nearest neighbors in the integral equation (3), and the analytical integration within the observation sphere with the dimensionless radius *R* = 2 for the line field.

In Fig. 5b, the BEM-FMM solution for every field component is shown by solid curves; the ANSYS Maxwell 3D FEM solution for the same field components is shown by dotted curves. We do not show the solution in air since ANSYS Maxwell becomes inaccurate in this case.

Table 7 summarizes values of the least squares difference between the solutions obtained via BEM-FMM and FEM, respectively, for all three electric-field components within the head, and for all ten head models. The relative difference percentage was computed along the line shown in Fig. 5. Results of Fig. 5 are marked grey. Note that the BEM-FMM solution runs approximately 1,000 times faster than ANSYS Maxwell 3D (including meshing time) on the same server. This is also faster than reported recently (Makarov et al, 2018); a further speed improvement has been achieved by using GMRES and converting native MATLAB loops in Eq. (6) to executable FORTAN DLLs running within the MATLAB shell.

**Table 7.** Least squares difference percentage between the BEM-FMM and the finite element method solutions for the three field components within the head, and for ten head models of the Population Head Model Repository. The relative difference percentage was computed along the line shown in Fig. 5. Results of Fig. 5 are marked grey.

| Head model # | Difference for $E_y$ (largest field component, %) | Difference for $E_z$ (secondary field component, %) | Difference for $E_x$ (vanishingly small field component, %) |
| --- | --- | --- | --- |
| 101309 | 3.5 | 13.1 | 28.0 |
| 103111 | 1.4 | 10.5 | 17.0 |
| 105014 | 3.2 | 9.2 | 55.5 |
| 105115 | 1.8 | 9.2 | 31.7 |
| 106016 | 1.8 | 6.8 | 34.8 |
| 110411 | 2.5 | 13.0 | 22.5 |
| 111716 | 1.6 | 10.6 | 27.8 |
| 113619 | 1.6 | 6.3 | 32.2 |
| 117122 | 3.1 | 18.7 | 27.9 |
| 118932 | 1.6 | 12.6 | 15.5 |

The excellent numerical agreement established in Table 7 for the dominant field component strongly supports the high accuracy of our method. To the authors’ knowledge, the present comparison scenario is by itself the first comprehensive example of this kind.

## 5. Discussion and Conclusion

Despite significant potential advantages quantified above, the BEM-FMM algorithm is not without its limitations. The FMM portion of the BEM-FMM algorithm is quite nontrivial in implementation. The BEM piece, on the other hand, at present relies upon tuning several parameters (number of neighbor integrals, terminating relative residual) in order to obtain a good convergence.

In contrast to this, the FEM algorithm has been extensively studied and applied across various engineering disciplines for decades; there are many highly reliable solvers available. Also, an application-specific implementation of FEM, coupled with novel solvers such as algebraic multigrid (AMG) preconditioners (Henson et al., 2002), can significantly speed up calculations when compared with the general FEM environment and a classic multipurpose solver, with no loss of accuracy or stability.

Another issue is that of accessibility. The present implementation of the BEM-FMM relies upon proprietary software (MATLAB^®^) while the transcranial brain stimulation modeling software SimNIBS is entirely open-source.

When evaluating the error of field calculations on real subjects, we must also take into account other key model parameters such as quality of the segmentation (Nielsen et al., 2018) and uncertainties in tissue conductivity values (Weise et al., 2015). These factors may cause simulation errors orders of magnitude larger than the numerical errors observed in the sphere models.

Finally yet importantly, it has been discussed in many sources (see Opitz et al., 2018) that the main limitation of the BEM formulation at present resides in its inability to model tissue anisotropies in a straightforward way.

## Supporting information

python script to run SimNIBS

## Acknowledgements

The authors wish to thank Dr. Leslie Greengard of the Courant Institute of Mathematical Sciences, New York, NY for useful remarks. The authors are thankful to Dr. Angel Peterchev of Duke University, Durham, NC for his constructive criticism. Dr. Axel Thielscher of Technical University of Denmark and the Danish Research Centre for Magnetic Resonance greatly helped in initiating and shaping this study. This work has been partially supported by the National Institutes of Health Grant R01MH111829, Novo Nordisk Fonden (grant Nr. NNF14OC0011413), and Lundbeckfonden (grant Nr. R118-A11308).

